# Prediction of gene essentiality using machine learning and genome-scale metabolic models

**DOI:** 10.1101/2022.03.31.486520

**Authors:** Lilli J. Freischem, Mauricio Barahona, Diego A. Oyarzún

**Affiliations:** School of Informatics, The University of Edinburgh, Edinburgh, UK; Department of Mathematics, Imperial College London, London, UK; School of Biological Sciences, The University of Edinburgh, Edinburgh, UK; The Alan Turing Institute, London, UK

**Keywords:** machine learning, gene essentiality, flux balance analysis, metabolic modelling, network science

## Abstract

The identification of essential genes, i.e. those that impair cell survival when deleted, requires large growth assays of knock-out strains. The complexity and cost of such experiments has triggered a growing interest in computational methods for gene essentiality prediction. In the case of metabolic genes, Flux Balance Analysis (FBA) is widely employed to predict essentiality under the assumption that cells maximize their growth rate. However, this approach implicitly assumes that knock-out strains optimize the same objectives as the wild-type, which excludes cases in which deletions cause large changes in cell physiology to meet other objectives for survival. Here we resolve this limitation with a novel machine learning approach that predicts essentiality directly from wild-type flux distributions. We first project the wild-type FBA solution onto a mass flow graph, a digraph with reactions as nodes and edge weights proportional to the mass transfer between reactions, and then train binary classifiers on the connectivity of graph nodes. We demonstrate the efficacy of this approach using the most complete metabolic model of *Escherichia coli*, achieving near state-of-the art prediction accuracy for essential genes. Our approach suggests that wild-type FBA solutions contain enough information to predict essentiality, without the need to assume optimality of deletion strains.

## I. INTRODUCTION

The identification of essential genes can reveal the minimal functional modules that allow an organism to survive. Gene essentiality is of paramount importance in biomedicine and biotechnology, e.g. for identifying therapeutic targets^1,2^ or improving yield in strains engineered for chemical production^3^. The identification of essential genes requires screening assays with high-throughput techniques such as RNA interference or CRISPR-based screens^4^. Due to the cost of such screens, there is substantial interest in computational methods that can rapidly explore the impact of gene deletions and complement current experimental efforts to determine gene essentiality. Such approaches typically employ machine learning in combination with various properties such as sequence homology and gene-function ontologies^5,6^.

In the case of metabolic genes, i.e. those that code for metabolic enzymes, the most popular approach for essentiality prediction is Flux Balance Analysis (FBA), which computes genome-wide metabolic flux distributions on the basis of an optimization principle^7^. This approach allows to rapidly simulate deletions and has shown promising results in some model prokaryotes; for example, the most complete FBA model of *Escherichia coli* predicts essential genes with up to 93% accuracy^8^.

Although various studies have reported encouraging results beyond the prokaryotic world^9^, the ability of FBA to predict gene essentiality is much more limited in eukaryotes and higher order organisms^10^. This limitation is partly due to the varied quality of current genomescale metabolic models, as well as the sensitivity of FBA predictions to the choice of objective function to be optimised. A common objective function is growth rate maximization^11^, but it is unclear if deletion strains still attempt to maximize growth, or if instead gene deletions alter cell physiology to meet other objectives for survival. Most recently, there is the growing realization that the integration of FBA and machine learning offers substantial promise to overcome some of the inherent limitations of genome-scale metabolic models^12^.

In this paper we present a machine learning approach to predict gene essentiality with minimal assumptions on the underlying optimality of the metabolic network. Crucially, our method does not require the computation of FBA solutions for deletion strains, but predicts essentiality directly from the wild-type flux distribution. The method works by first projecting the flux distribution onto a mass flow graph^13^, i.e. a directed graph with reactions as nodes and edge weights as chemical mass flows, and then using the connectivity of graph nodes to train machine learning predictors of gene essentiality. Using data from a large growth assay in *Escherichia coli* ^8^, we trained a range of binary classifiers and demonstrate that our method allows to predict essential genes with near state-of-the-art accuracy. We conclude with a discussion of limitations of the method and directions for future improvement.

## II. PRELIMINARIES

### A. Flux balance analysis

Flux balance analysis (FBA) is a widely-adopted approach to study cellular metabolism^7^. Assuming that a metabolic network with *n* metabolites and *m* reactions is in steady state, the vector of reaction fluxes **v** satisfies the mass balance equation:

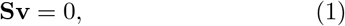

where **S** is an *n* × *m* integer matrix, and its *S*_*ij*_ entry corresponds to the net number of *X*_*i*_ molecules produced (*S*_*ij*_ *>* 0) or consumed (*S*_*ij*_ *<* 0) by reaction *R*_*j*_. In its simplest form, FBA finds the solution vector **v**^*∗*^ to the following constrained optimization problem:

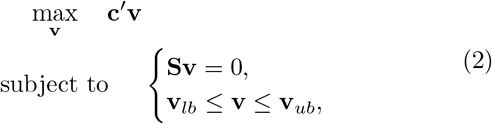

where **c** is a vector of flux weights, and (**v**_*lb*_, **v**_*ub*_) are lower and upper bounds on reaction fluxes, respectively. A common use case is to employ the flux bounds for predicting flux distributions in various genetic backgrounds or environmental conditions. For example, deletions of the gene encoding for reaction *v*_*j*_ can be simulated by setting *v*_*lb, j*_ = *v*_*ub, j*_ = 0, while specific growth conditions can be modelled by manipulating the bounds of nutrient uptake reactions.

### B. Construction of mass flow graphs

Mass flow graphs (MFGs) were originally introduced by^13^ as a method to map flux vectors onto a directed graph that can be analysed with tools from network science. In these graphs, nodes correspond to metabolic reactions and two nodes are connected if they share metabolites either as reactants or products. A key advantage of such reaction-centric graphs is that they avoid the need to prune pool metabolites, i.e. enzymatic cofactors, ions and others, that appear in many metabolic reactions. In other graph constructions, pool metabolites are manually removed so as to avoid the spurious connections produced by their high connectivity, which tend to dominate the topology of the resulting graph. In the MFG construction, pool metabolites are kept as-is and they map onto weak connections between graph nodes, thus reducing their impact on the overall connectivity.

To construct an MFG, we define the weight of the connection between reactions *R*_*i*_ and *R*_*j*_ as the total flux of metabolites produced by *R*_*i*_ that are consumed by *R*_*j*_ (Figure 1A). Mathematically, the adjacency matrix for an MFG can be directly constructed from the stoichiometric matrix **S** and any FBA solution vector **v**^*∗*^. We first unfold the flux vector into 2*m* forward and reverse reactions:

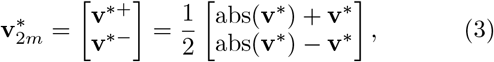

and define the corresponding stoichiometric matrix as:

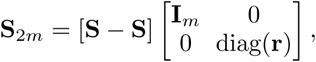

where **r** is the *m*-dimensional reversibility vector such that

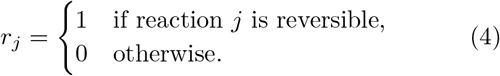

**FIG. 1.**
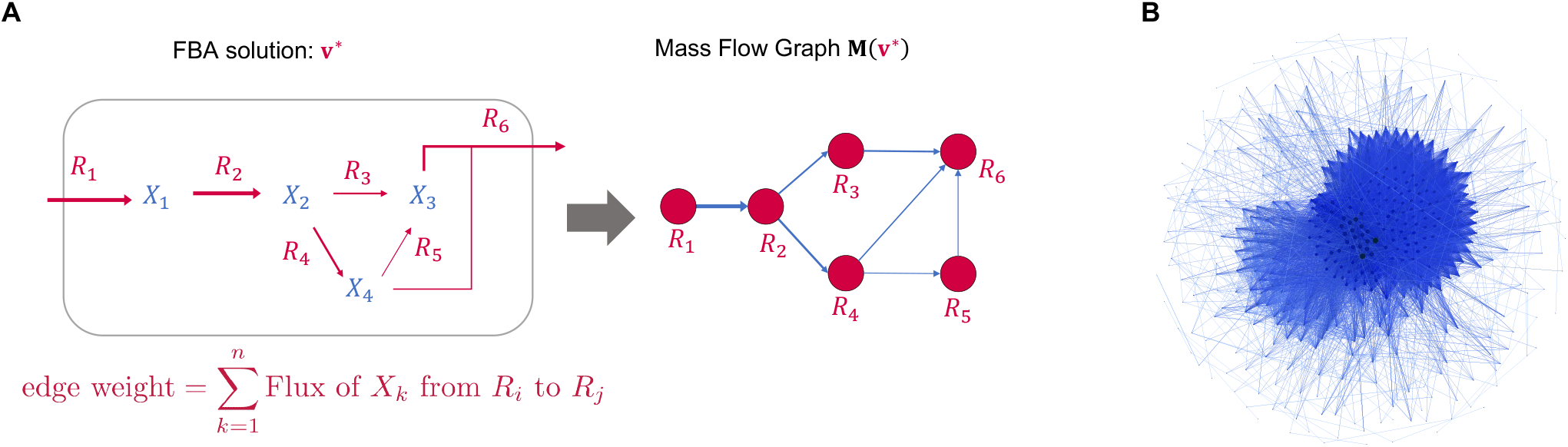
Construction of mass flow graphs. **(A)** Mass flow graphs (MFG) can be directly built from the stoichiometric matrix and a flux vector^13^. The MFG is a directed graph with reactions as nodes, and edge weights corresponding to the total mass flow between two reactions, defined by the adjacency matrix in Eq. (5). **(B)** Mass flow graph of *Escherichia coli* under aerobic growth with glucose as sole carbon source; the graph contains *k* = 444 reaction nodes and 14,459 edges, and was computed from the most complete genome-scale reconstruction (iML1515) by^8^.

The adjacency matrix of the MFG is then given by:

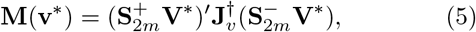

where † denotes the matrix pseudoinverse, and the matrices are defined as 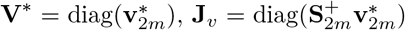, with

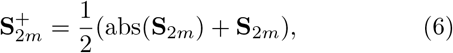

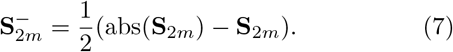

As explained in detail in^13^, the edge weights in the adjacency matrix **M**(**v**^*∗*^) have units of mass per unit of time, and represent the strength of connectivity between reactions in terms of the chemical mass flow they share. As an illustration, Figure 1B shows the MFG for the most complete metabolic reconstruction of *Escherichia coli* iML1515 described in^8^, for cells growing aerobically in glucose; Table I shows a summary of the reaction flux bounds.

**TABLE I.** Reaction flux bounds of *E. coli* model iML1515. Values were modified to simulate aerobic growth using glucose as sole carbon source. The resulting MFG is shown in Figure 1B.

| Reactions | Bounds |
| --- | --- |
| L-arabinose isomerase (ARAI),<br>L-ribulokinase (RBK_L1),<br>Rhamnulose-1-phosphate aldolase (RMPA),<br>Lyxose isomerase (LYXI),<br>L-rhamnose isomerase (RMI),<br>Rhamnulokinase (RMK), and<br>B-galactosidase (LACZ) | $\mathbf{v}_{lb} = \mathbf{v}_{ub} = 0$ |
| Oxygen exchange (EX_o2_e) | $\mathbf{v}_{lb} = -20$ |
| Glucose uptake (EX_glc_D_e) | $\mathbf{v}_{lb} = -10$ |

### C. Binary classification

Binary classification is one of the core problems in supervised machine learning. The goal is to extract patterns from observations of two classes of objects, and use these patterns to automatically determine the class of a new, unseen, object. In our case, we are interested in using growth measurements from knock-out screens to build automated classifiers that determine whether a gene is essential or non-essential. More specifically, assume we have *N* pairs:

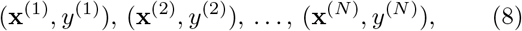

where **x**^(*i*)^ ∈ ℝ^*p*^ is a *p*-dimensional vector of *features* associated with the *i*^th^ gene, and *y*^(*i*)^ ∈ {0, 1} is the *label* or *class* of the *i*^th^ gene. Without loss of generality, we denote non-essential and essential genes as the negative class (0) and positive class (1), respectively.

The feature vectors and labels are assembled into a feature matrix **X** and a vector of class labels **y**:

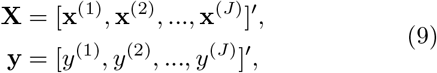

that are employed to train a classification algorithm to learn the patterns in **X** required to predict the labels in **y**. Once trained, the algorithm can automatically assign labels to new samples on the basis of their feature vectors. Given the feature vector of a new sample 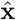, the binary classifier predicts the corresponding class label *ŷ* based on the learned classification rules.

There are various types of classification models, typically based on different assumptions on the shape of the feature space. Common model include logistic regression, decision trees, neural networks, and support vector machines^14^. The choice of specific models depends largely on the task and dataset at hand, and in a typical machine learning pipeline model training is accompanied by cross-validation analyses to prevent overfitting and perform model selection.

## III. BINARY CLASSIFIERS TRAINED ON THE MASS FLOW GRAPHS

We built a machine learning pipeline (Figure 2) to train binary classifiers that predict the essentiality labels from features extracted from the mass flow graphs. Next we detail each step of our approach.

**FIG. 2.**
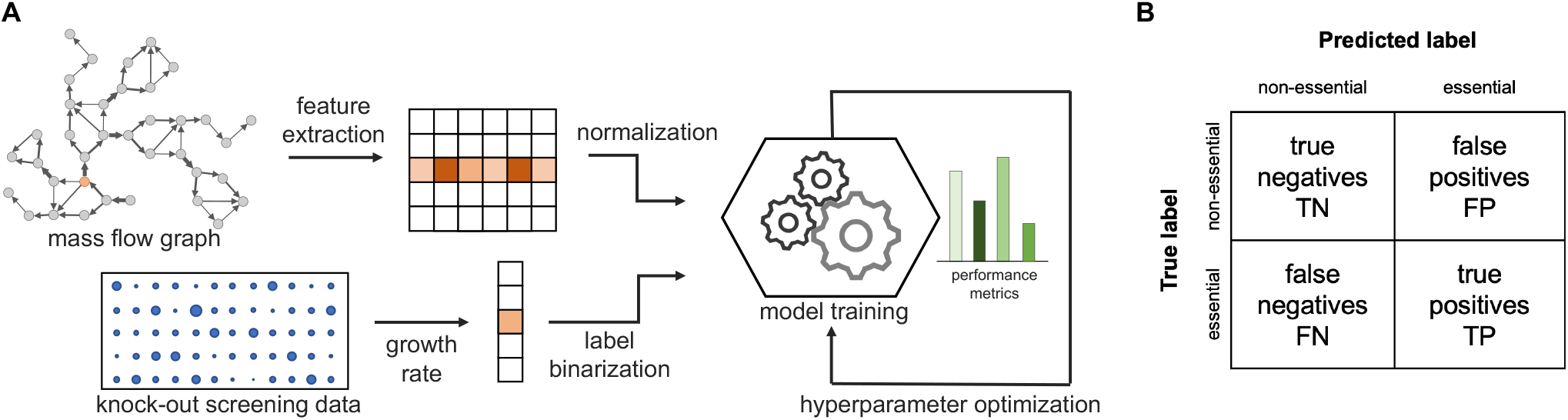
Training binary classification algorithms on mass flow graphs. (**A**) The node feature matrix **X** is computed from the adjacency matrix of a mass flow graph **M**. Binary classifiers are trained using **X** and measured essentiality labels **y**. For hyperparameter optimization, different performance metrics were employed to account for the imbalanced number of essential and non-essential genes.(**B**) Confusion matrix to assess the performance of a binary classifier.

### A. Feature extraction

We first featurize each gene using the connectivity of the corresponding reaction node in the MFG. To construct a feature matrix from the adjacency matrix **M** of the MFG, we note that FBA solution vectors tend to be sparse, i.e. they contain many reactions that carry zero flux. Such reactions map onto disconnected nodes in the MFG, and thus correspond to non-essential genes. If the *i*^th^ node is disconnected, the corresponding *i*^th^ row and column in **M** contain only zeros. We removed such disconnected nodes, so that if the connected component of the MFG contains *k* nodes, we compute a reduced adjancency matrix **M**_*k*_ where all (*m* − *k*) zero rows and columns have been removed. We then define a feature matrix as

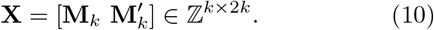

The feature matrix has been augmented to include the weights of incoming and outgoing edges as features for each reaction node. In other words, the *i*^th^ row of **X** contains both the outgoing and incoming edge weights of reaction *R*_*j*_ in the MFG. We also note that the feature matrix tends to be sparse, because reactions typically share a reduced number of metabolites and thus feature vectors have a large number of zero entries.

### B. Data normalization and labelling

Many classification algorithms are based on distances between feature vectors that can be sensitive to scaling; the Euclidean or cosine distances are classic examples of such scale sensitive distance functions. To avoid problems with feature scaling, we normalized the feature matrix **X** prior to model training. This is usually achieved by subtracting the feature mean and scaling to unit variance. However, in our case we only scale features to unit variance so as to preserve the sparsity structure of the feature matrix:

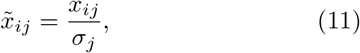

where 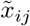 is the normalized entry of the feature matrix **X**, and *σ*_*j*_ is the standard deviation of feature *j*. The normalization factors are computed from the training data and stored to transform future inputs accordingly.

To obtain essentiality labels for each gene, we employ growth assay data that quantify the impact of a single deletion on cellular growth. If *g*_*W T*_ is the measured growth rate of the wild-type, and *g*_*i*_ is the growth rate of a strain with the *i*^th^ gene knocked-out, the essentiality score is:

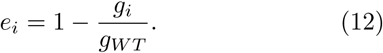

In practice, most genes have essentiality scores close to 0 or 1, so we binarize them to obtain the class label for each gene:

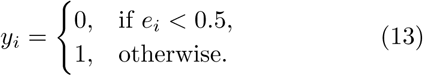

### C. Model training and hyperparameter optimization

We considered a number of classification algorithms, including logistic regression, support vector machines, neural networks and ensemble classifiers based on decision trees. These models were trained using optimization routines available in the scikit-learn Python package, using suitably defined loss functions representing the quality of the classification performance; the loss function employed depends on the model under consideration, and common examples include cross-entropy loss, logistic loss and hinge loss^15^. All models were trained on a fixed fraction of the *k* connected nodes of the MFG, and we held out a subset of the graph nodes to test model performance on unseen data; the data were split using stratified sampling to account for the imbalance between the number of essential and non-essential reactions. From the classification results on the test set, we compute the confusion matrix illustrated in Figure 2B to compare performance across different models. Performance was quantified using five classification scores:

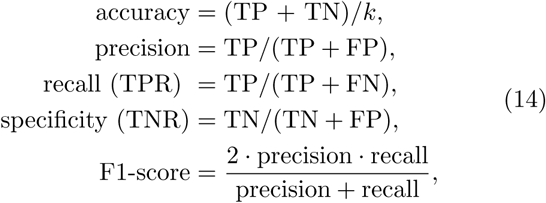

which are calculated from the number of true positives (TP), true negatives (TN), false positives (FP) and false negatives (FN) in the confusion matrix (Figure 2B).

In addition to the learned parameters, each classification algorithm has a number of hyperparameters that define various properties and settings of the model. In a typical machine learning pipeline, the hyperparameters are not fitted directly to data, but instead employed to perform model selection or decide between several competing classifiers according to a performance metric. The process of choosing suitable hyperparameters can quickly become computationally expensive since it requires training the model for each combination of hyperparameters across a large grid. We instead employed Bayesian optimization^16^, a global, gradient-free, optimization method purposely designed to reduce computational costs in problems with expensive objective functions. In this approach, the objective function is assumed to be a random variable. After a number of function evaluations, a prior on the objective function is updated to compute a posterior distribution over the objective function, typically using a Gaussian Process regressor. The posterior is then employed to determine the next combinations of hyperparameters for model training using a suitable acquisition function that balances exploration and exploitation of the hyperparameter space. The Bayesian optimization strategy provides an efficient alternative to methods based on grid search or gradient ascent.

## IV. APPLICATION TO ESCHERICHIA COLI METABOLIC NETWORK

To illustrate our approach, we applied our pipeline to predict gene essentiality using the metabolic reconstruction iML1515 of *E. coli* MG1655 strain, described by^8^. That work also provides growth assay data for strain BW25113 that can be compared with predictions from our machine learning algorithm.

We computed the MFG of the iML1515 model assuming aerobic growth with glucose as sole carbon source. Since *E. coli* BW25113 lacks several genes from MG1655, we set flux bounds of the associated reactions to zero. Additionally, we adjusted the oxygen exchange reaction bounds to simulate aerobic growth and the glucose uptake reaction bounds as glucose was used as primary carbon source. The employed flux bounds are summarised in Table I.

Since growth assay data contains deletions of genes, not reactions, the essentiality labels must be converted from the space of genes onto the space of reactions. To this end, we employed the gene-protein-reaction (GPR) boolean rules included in the iML1515 genome-scale model. These rules describe the dependency of metabolic reactions on metabolic genes included in the genomescale model. We found that iML1515 contains only 155 reactions that map one-to-one from genes to reactions (Figure 3A). To resolve this limitation and expand the number of reactions for model training, we also included reactions that are deactivated upon deletion of a specific gene, even if that deletion also deactivates other reactions. Under this definition, we could assign essentiality labels to a total of 255 reactions in the MFG, which amounts to a coverage of ∼57% of all active reactions (Figure 3A). Reactions that cannot be deactivated by a single knockout were excluded from model training because their essentiality labels cannot be inferred from the available growth data.

**FIG. 3.**
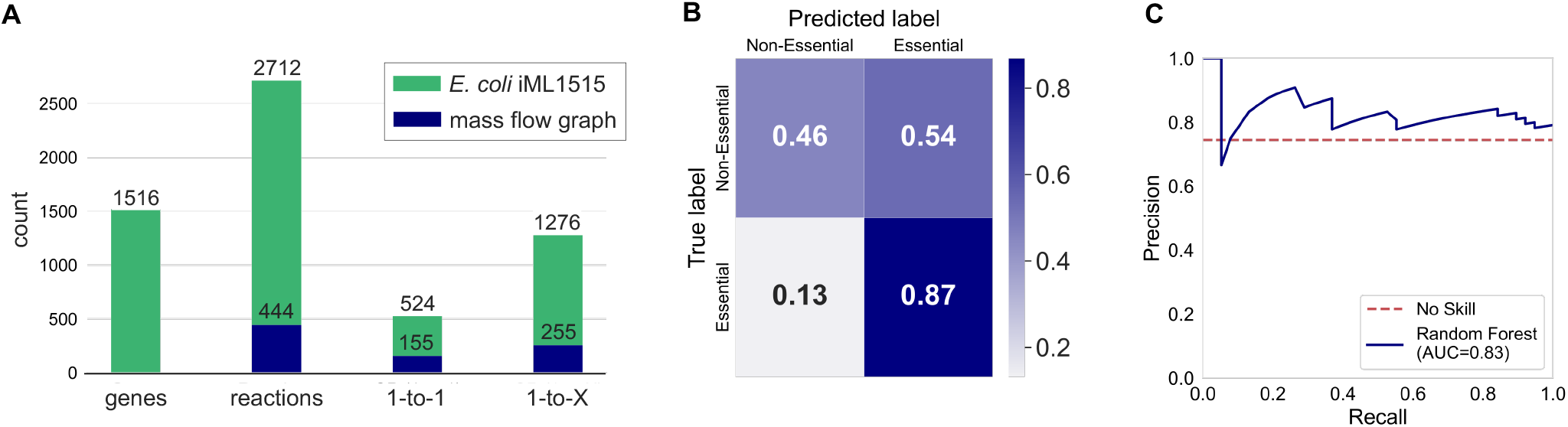
Prediction of gene essentiality in *Escherichia coli* iML1515. (**A**) Reaction and gene counts in iML1515. Shown are the total number of genes, total number of reactions, and the number of reactions that are deactivated by single knockouts that do not deactivate any other reactions (“1-to-1”) and that deactivate one or more reactions (“1-to-X”). (**B**) Normalized confusion matrix of the best performing model: a random forest classifier with hyperparameters tuned to maximize the macro-averaged F1-score. (**C**) Precision-recall curve of the Random Forest model. Panels B and C were computed on a test set with 20% of graph nodes. We note that an unskilled classifier has a precision of ∼75%, which corresponds to the class imbalance between essential and non essential genes.

We first trained four different models for binary classification and used 5-fold cross-validation to optimize hyperparameters, compare model performance, and check for overfitting. Hyperparameters were determined with Bayesian optimization using the macro-averaged F1score as objective function (*F*_1, macro_); this score is the arithmetic mean of the F1-scores computed for essential and non-essential genes separately, as otherwise the imbalance between both classes can cause the optimizer to converge to classifiers that are unable to classify the minority class (non-essential reactions in our case). The results in Table II summarize the performance of models with optimized hyperparameters when trained on 80% of reactions and tested on a held-out set with the remaining 20% of reactions.

**TABLE 2.** Classification performance of four binary classifiers. Results show performance metrics for Random Forest (RF), Multi-Layer Perceptron (MLP), Logistic Regression (LogReg), and C-Support Vector Machine (SVC). Models were trained on 80% of the nodes of the MFG; the reported performance metrics were computed on a held-out dataset with 20% of nodes.

| model | accuracy | precision | recall | F1 | $F_{1, \text{macro}}$ |
| --- | --- | --- | --- | --- | --- |
| RF | 0.76 | 0.82 | 0.87 | 0.85 | 0.67 |
| MLP | 0.73 | 0.80 | 0.84 | 0.82 | 0.62 |
| LogReg | 0.67 | 0.76 | 0.82 | 0.78 | 0.52 |
| SVC | 0.61 | 0.74 | 0.74 | 0.74 | 0.48 |

We found the best model to be a Random Forest classifier consisting of 300 trees with a maximum depth of 50, and using information gain as criterion and log_2_(2*k*) features to determine optimal tree splits. In crossvalidation, the RF model had 82% precision, 86% recall and an F1-score of 83%. Evaluated on the test set, the model had an overall accuracy of 76% with a precision of 82.5% and a recall of 86.8% (see Figure 3B–C). For comparison, the FBA predictions^8^ on the same set of genes have 84.3% accuracy, and both precision and recall of 89.5%. Despite the small amount of data available to train our classifiers, our predictions are promisingly close to those given by FBA; since our approach does not assume optimality of the knockout strains, these results suggest that wild-type flux distributions may contain sufficient information to predict the essentiality of knockouts. Examination of the confusion matrix (Figure 3B) suggests that our classifier is comparatively poor at predicting the non-essential reactions, but shows near state-of-the-art accuracy for essential genes.

## V. CONCLUSION

In this paper we have described a new computational method for predicting essentiality of metabolic genes. Our approach combines flux balance analysis (FBA) with machine learning algorithms trained on growth assay data. By projecting flux vectors onto a directed graph, we recast the problem as a binary classification task on the graph nodes. Test results on a reduced gene set in *Escherichia coli* suggest that the method can deliver predictions that are close to those delivered by FBA. Unlike such methods, however, our approach does not require the computation of FBA solutions for deletion strains and relies solely on the wild-type flux distribution. This is important because it assumes optimality of the wildtype, but not of the deletion strains, and thus can account for scenarios in which deletions cause changes in cellular objectives.

Despite an increased interest in machine learning for essentiality prediction, current methods still suffer from limitations in their accuracy and ability to generalize across environmental conditions or species^6^. This is partly due to the lack of gene featurization strategies that are predictive of essentiality. Here we have employed graph connectivity as features to train our models, and our results show some promise for their wider applicability in other organisms.

Our aim in this paper was to provide the methodological foundations for a new strategy that combines elements of machine learning and flux balance analysis, two of the leading approaches in the field. There are several important points for improvement that deserve further exploration. For example, we have only tested the method in a single growth condition for *Escherichia coli* (aerobic growth in glucose), and further tests should be carried out to establish the validity of the method across other conditions. This is particularly relevant for conditionally essential genes, i.e. those on which survival depends on the specific environmental conditions. We also observed comparatively poor prediction accuracy for non-essential genes; this is likely because these tend to be underrepresented in FBA solutions, which produces imbalanced datasets for the binary classification problem. Solutions to this caveat will likely require strategies to mitigate the impact of imbalanced datasets, such as under- and over-sampling, or the use of different penalties for misclassification errors in the loss function.

The identification of essential genes is of fundamental importance in basic science, biomedicine, biotechnology and a number of other disciplines. In this paper, we developed a novel method for essentiality prediction which we hope will prove a catalyst for further exploration of machine learning methods in the field.

## APPENDIX

## List of genes in the test set

We evaluated the performance of our machine learning algorithm on a test set with 51 genes, corresponding to 20% of all nodes in the MFG. All these genes were held out when training the models in Table II: *adk, argA, argH, aroB, bioB, bioF, dapB, dapF, deoB, dxr, eno, fabG, fabZ, fadE, folE, glmM, glmS, glmU, gltX, glyA, gmk, gnd, gsk, hemD, hisA, hisD, iscU, ispU, lpxC, ltaE, murC, nadB, nadC, panD, pssA, purA, purC, purM, purN, ribE, serA, tesB, thiE, thiG, thiL, tnaA, trpD, ubiD, waaA, yrbG, zupT*.

